## Supplementary Figures and Tables for "Specific length and structure rather than high thermodynamic stability enable regulatory mRNA stem-loops to pause translation"

### Supplementary Information

### Supplementary Figures

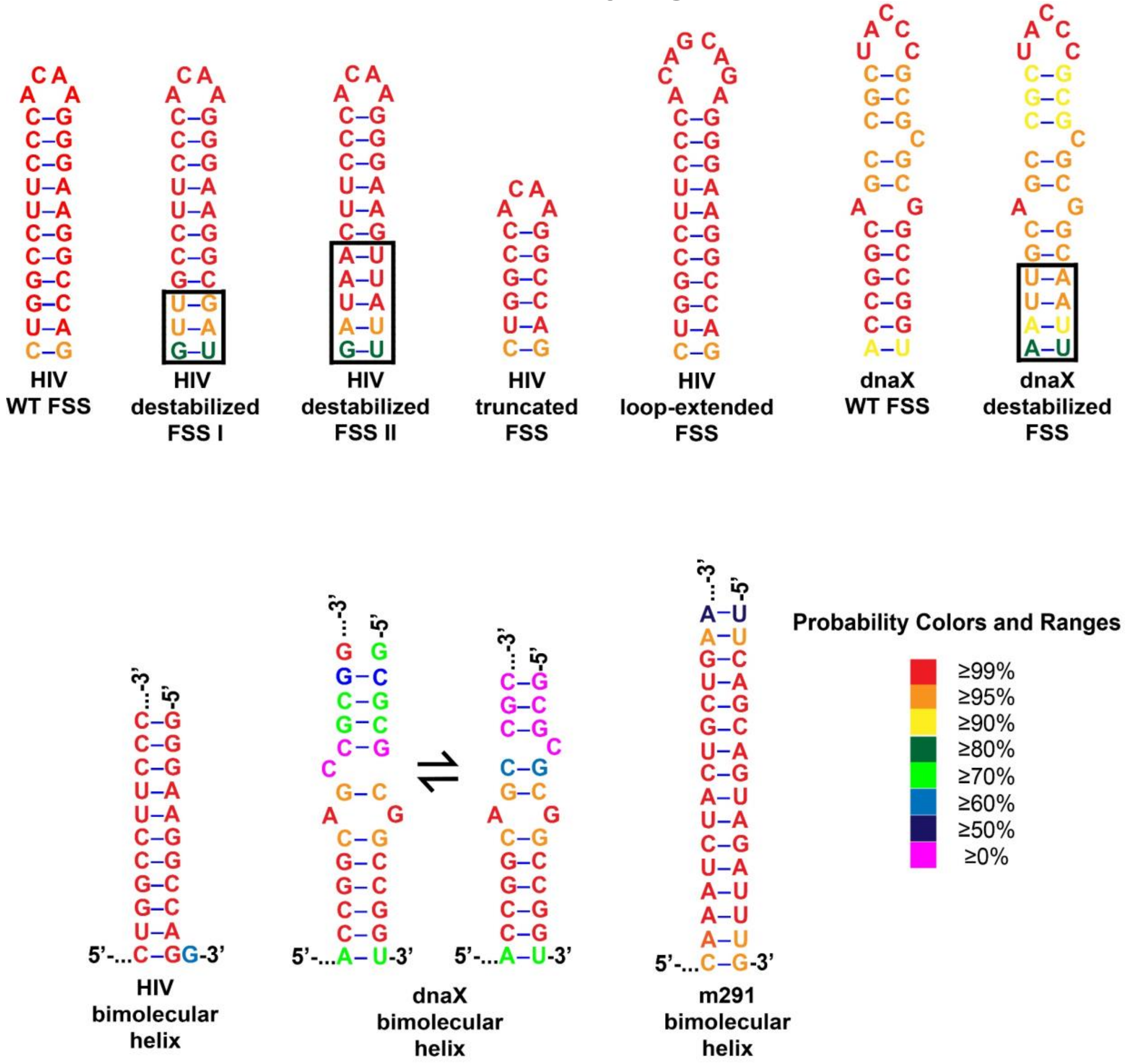


**Supplementary Figure 1. Basepair probabilities for model mRNAs (shown in Fig. 1).** Maximum expected accuracy structures were calculated using RNAstructure[1], and estimated base pairing probabilities are color annotated. The hairpin stem-loop sequences were mutated to have altered stability, but similarly high pairing probabilities as the wild-type sequences. The dnaX bimolecular helix is predicted to have two structures in equilibrium with folding free energy changes of is -20.3 kcal/mol (left) and -20.2 kcal/mol (right). The conformational difference is whether the C bulges in the mRNA (left) or the trans oligonucleotide (right; more similar to the dnaX hairpin structure). The base of the helix has similarly high probability as the dnaX wild-type hairpin stem-loop.

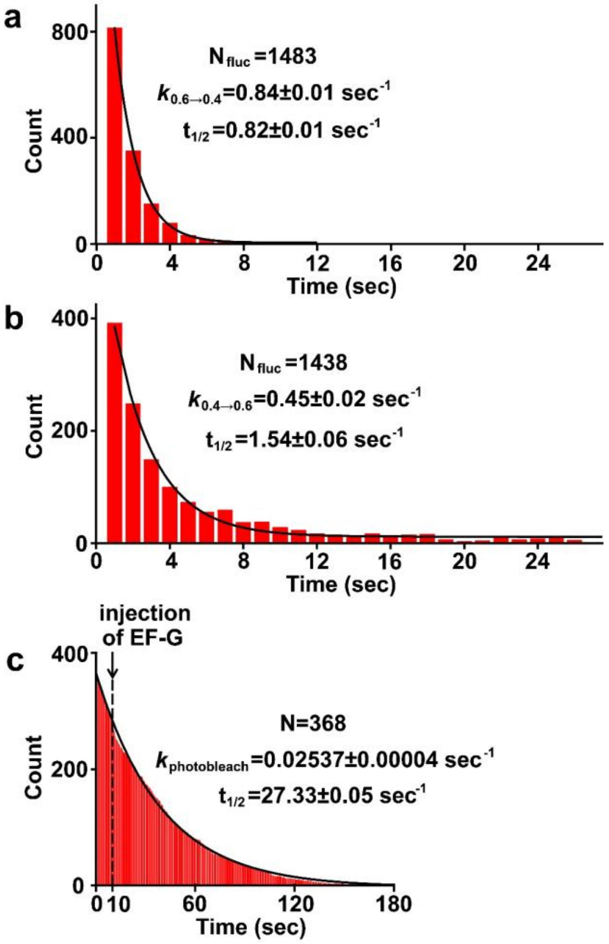


**Supplementary Figure 2. Spontaneous fluctuation and photobleaching rates of the S6-cy5/L9-cy3 ribosomes.**

**(a-b)** Histograms (1 s binning size) show dwell time of the NR (0.6 FRET) **(a)** and R (0.4 FRET) **(b)** states of the S6/L9-labeled 70S ribosomes containing deacylated $tRNA^{Phe}$ in the P site. Transition rates and half-lives of dwell time were deduced from single exponential fitting shown by black curves. **(c)** Histogram (0.1 s binning size) show distribution of cy5 life times in translocation smFRET measurments with continues Cy3 excitation in ribosomes programmed with “wild-type” HIV FSS and dnaX FSS mRNAs. Injection of EF-G•GTP is indicated by the arrow. Photobleaching rate and half-life were deduced from single exponential fitting shown by black curve. N indicates the number of FRET traces incorporated into each histogram.

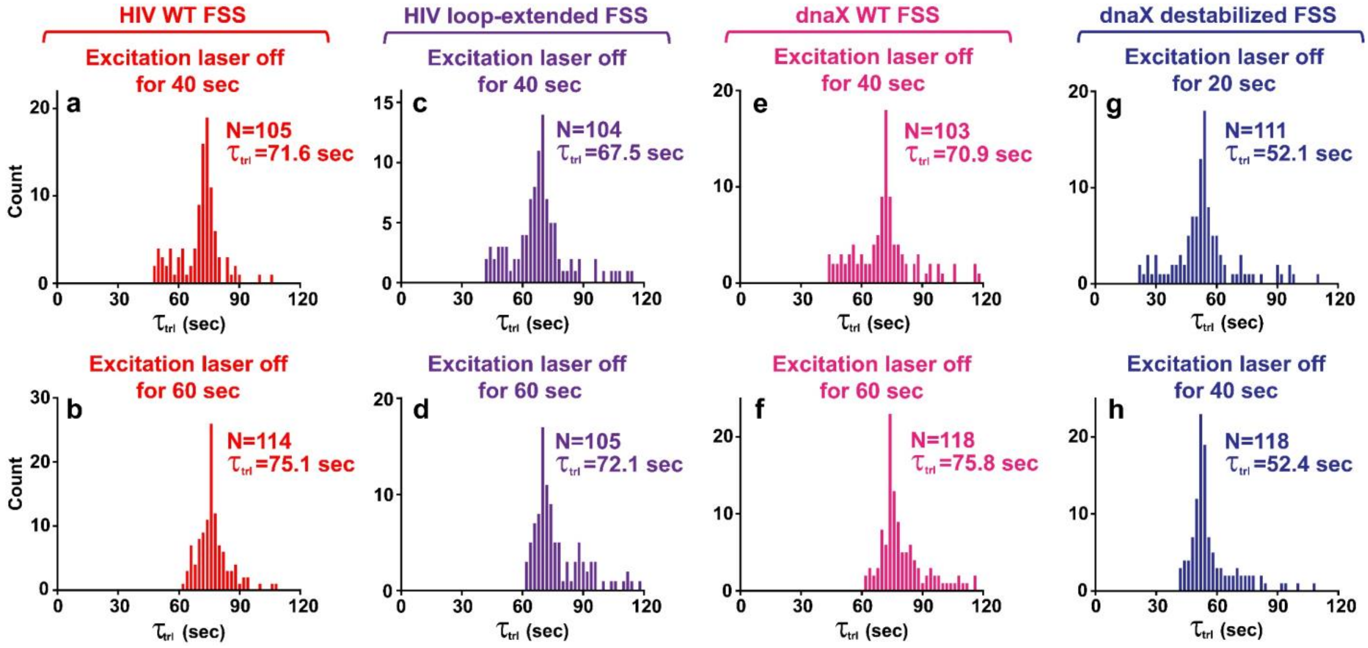


**Supplementary Figure 3. Variations of the interval, during which the excitation laser was switched off, did not affect distributions of $\tau_{trl}$.**

Kinetics of translocation was measured by smFRET experiments with pre-translocation S6-cy5/L9-cy3 ribosomes programmed mRNA containing HIV WT FSS (**a, b**), HIV loop-extended FSS (**c, d**), dnaX WT FSS (**e, f**) or dnaX destabilized FSS (**g, h**), which was positioned 11 nucleotides downstream of P-site codon. Because these FSS variants strongly inhibited translocation, the excitation laser was turned off after the EF-G injection and switched back on either 20, 40 or 60 s later to extend lifetime of the acceptor fluorophore as indicated. Histograms (2 s binning size) show distributions and median values of $\tau_{trl}$.

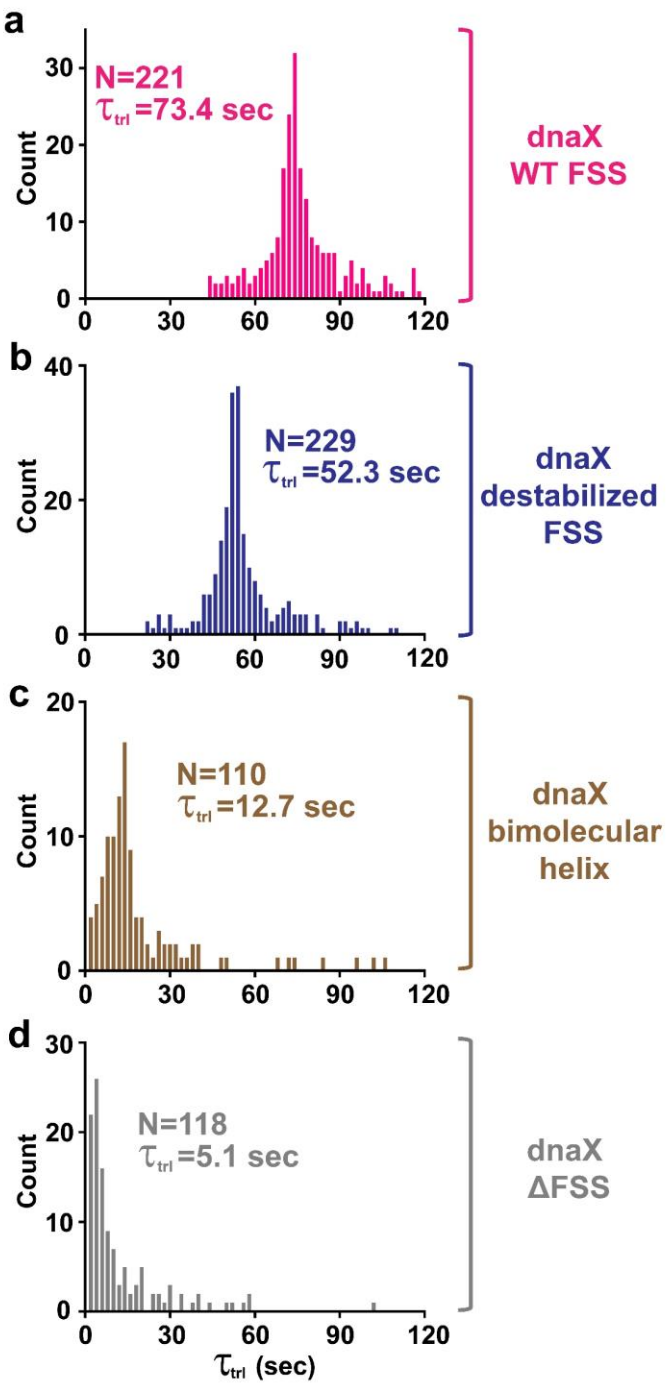


**Supplementary Figure 4. The dnaX FSS-induced inhibition of translocation is alleviated by elimination of the loop.** Kinetics of translocation was measured by smFRET experiments with pre-translocation S6-cy5/L9-cy3 ribosome programmed with dnaX WT FSS mRNA **(a)**, dnaX destabilized FSS mRNA **(b)**, dnaX bimolecular helix mRNA **(c)** or dnaX ΔFSS mRNA **(d)**. In each mRNA, the spacer between P-site codon and the downstream secondary structure was 11 nucleotides long. Histograms (2 s binning size) show the distributions and median values of $\tau_{trl}$. N indicates the number of FRET traces incorporated into each histogram.

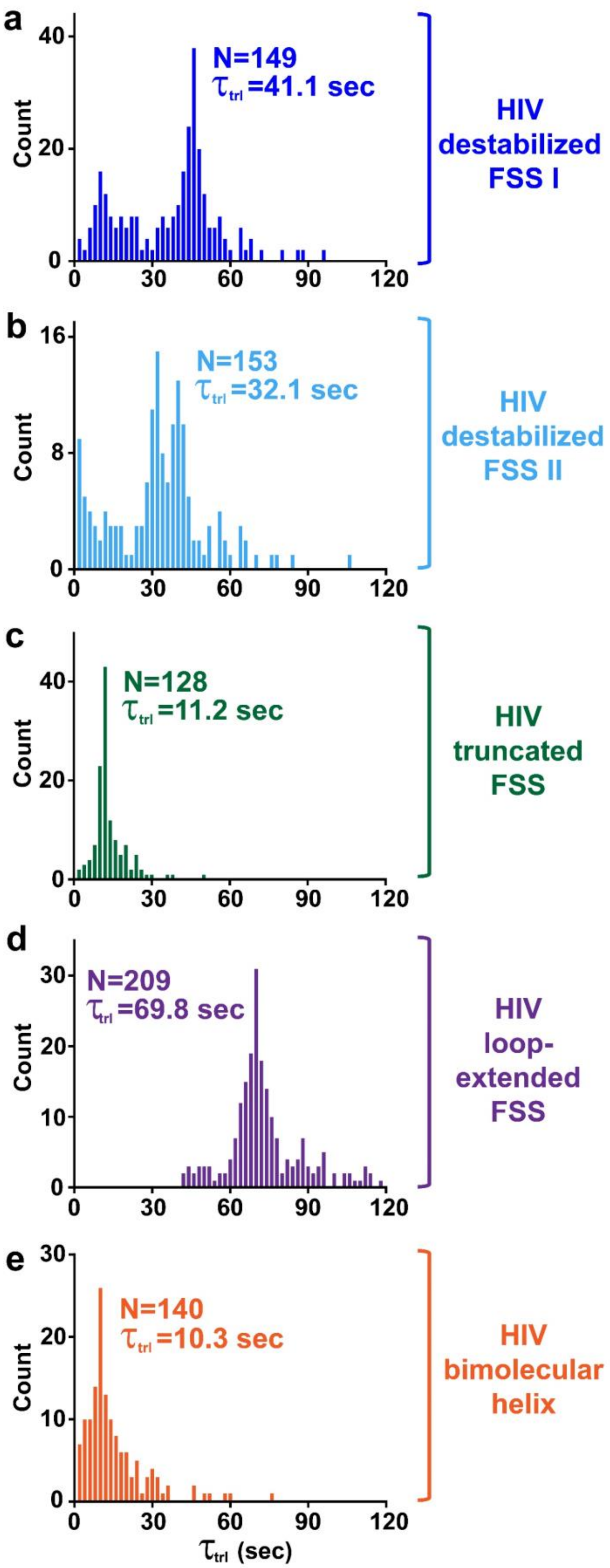


**Supplementary Figure 5. The HIV FSS-induced inhibition of translocation is alleviated by truncation of the stem or elimination of the loop.**

Kinetics of translocation was measured by smFRET experiments (**Fig. 2b**) with pre-translocation S6-cy5/L9-cy3 ribosome programmed with mRNAs containing HIV destabilized FSS I (**a**), HIV destabilized FSS II (**b**), HIV truncated FSS (**c**), HIV loop-extended FSS (**d**) or the bimolecular helix mimicking the native FSS stem (**e**). In each mRNA, the spacer between P-site codon and the downstream secondary structure was 11 nucleotides long. Histograms (2 s binning size) show the distributions and median values of $\tau_{trl}$. N indicates the number of FRET traces included in each histogram.

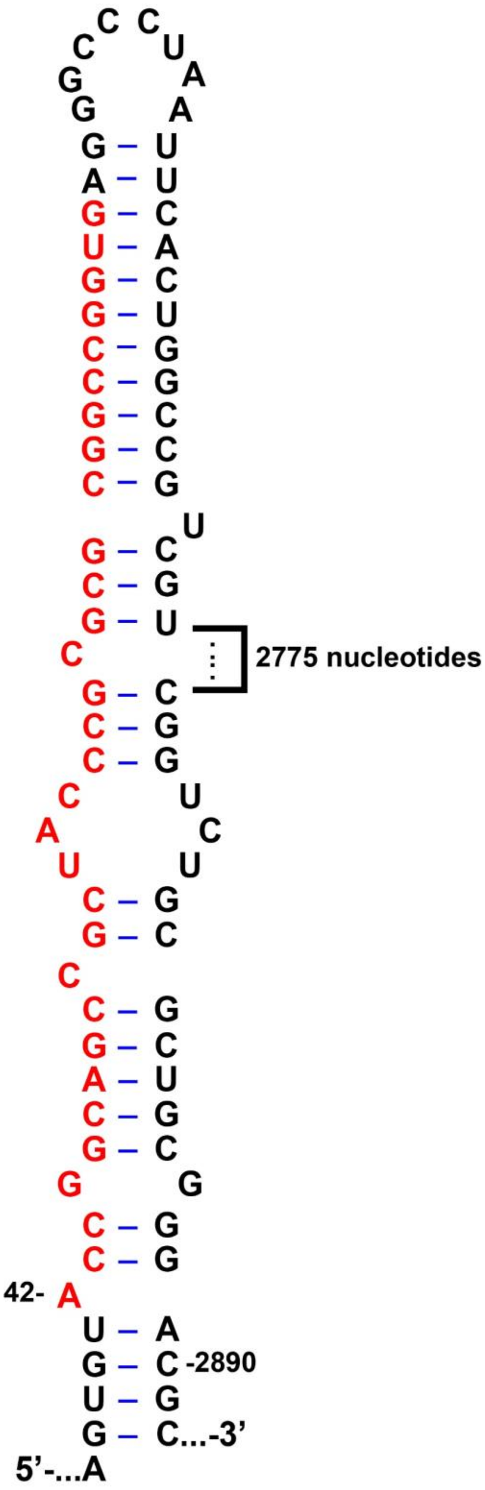


**Supplementary Figure 6. The alternative structure to the dnaX hairpin in the Atkins et al[2]. frameshifting assay.** This alternative structure is the predicted maximum expected accuracy structure, using RNAstructure, where the dnaX hairpin nucleotides are colored red. This alternative structure demonstrates basepairs between the dnaX sequence and nucleotides just 3’ downstream to the dnaX sequence and also nucleotides at the 3’ end of the β-galactosidase open reading frame (ORF). These pairs would compete with the dnaX hairpin stem-loop basepairs, complicating the interpretation of mutations in the frameshifting element. The sequence of the dnaX reporter was reconstructed based on the β-galactosidase ORF sequence and fragments of the reporter sequence shown in Fig. 2 of Larsen at el 1994[3] and Fig. 1 of Larsen et al 1997[2].

**Supplementary table 1. RNA sequences**

| RNA identifier | RNA sequence (5' to 3') |
|---|---|
| HIV_NS ΔFSS mRNA | **GGUUUUUCUUCUGAAGAUAAAG**CAACAACAACAAGGCAAGGAGGUAAAAAUGUUCUACAA |
| HIV_NS **WT FSS** mRNA (11-nt spacer) | **GGUUUUUCUUCUGAAGAUAAAG**CAACAACAACAAGGCAAGGAGGUAAAAAUGUUCUACAAGAU**CUGGCCUUCCCACAAGGGAAGGCCAG**GGAA |
| HIV_NS **WT FSS** mRNA (12-nt spacer) | **GGUUUUUCUUCUGAAGAUAAAG**CAACAACAACAAGGCAAGGAGGUAAAAAUGUUCUACAAGAAU**CUGGCCUUCCCACAAGGGAAGGCCAG**GGAA |
| HIV_NS **WT FSS** mRNA (13-nt spacer) | **GGUUUUUCUUCUGAAGAUAAAG**CAACAACAACAAGGCAAGGAGGUAAAAAUGUUCUACGGAAGAU**CUGGCCUUCCCACAAGGGAAGGCCAG**GGAA |
| HIV_NS **truncated FSS** mRNA | **GGUUUUUCUUCUGAAGAUAAAG**CAACAACAACAAGGCAAGGAGGUAAAAAUGUUCUACAAGAU**CUGGCCACAAGGCCAG**GGAA |
| HIV_NS **destabilized FSS I** mRNA | **GGUUUUUCUUCUGAAGAUAAAG**CAACAACAACAAGGCAAGGAGGUAAAAAUGUUCUACAAGAU**GUUGCCUUCCCACAAGGGAAGGCGAU**GGAA |
| HIV_NS **destabilized FSS II** mRNA | **GGUUUUUCUUCUGAAGAUAAAG**CAACAACAACAAGGCAAGGAGGUAAAAAUGUUCUACAAGAU**GAUAACUUCCCACAAGGGAAGUUAUU**GGAA |
| HIV **bimolecular helix** RNAs | **GGUUUUUCUUCUGAAGAUAAAG**CAACAACAACAAGGCAAGGAGGUAAAAAUGUUCUACAAGAU**CUGGCCUUCCC**GGAA<br>**GGGAAGGCCAGG** |
| dnaX_NS ΔFSS mRNA | **GGUUUUUCUUCUGAAGAUAAAG**CAACAACAACAAGGCAAAGGGAGCAACCAUGGUAUUCUACAGAGAACCGG |
| dnaX_NS **WT FSS** mRNA | **GGUUUUUCUUCUGAAGAUAAAG**CAACAACAACAAGGCAAAGGGAGCAACCAUGGUAUUCUACAGAGA**ACCGGCAGCCGCUACCCGCGCGCGGCCGGU**GAAUAACGGGAUC |
| dnaX_NS **destabilized FSS** mRNA | **GGUUUUUCUUCUGAAGAUAAAG**CAACAACAACAAGGCAAAGGGAGCAACCAUGGUAUUCUACAGAGA**AAUUGCAGCCGCUACCCGCGCGCGGCAAUU**GAAUAACGGGAUC |
| dnaX **bimolecular helix** RNAs | **GGUUUUUCUUCUGAAGAUAAAG**CAACAACAACAAGGCAAAGGGAGCAACCAUGGUAUUCUACAGAGA**ACCGGCAGCCGC**GGAUC<br>**GCGCGCGGCCGGU** |
| m291 **bimolecular helix** RNAs | GUAAAGUGUCAUAGCACCAACUGUUAAUUAAAUUAAAUUAAAAAGGAAAUAAAAAUGUUUGUAUACAAAUCUACUGCUGAACUCGCUGCACAAAUGGCUAAACUGAAUGGCAAUAAA**GGUUUUUCUUCUGAAGAUAAAG**<br>**UUCAGCAGUAGAUUUG** |

The RNA sequences are shown in 5'-to-3' direction. In each RNA, the Shine-Dalgarno-like sequence and the RNA spacer extending from the first nucleotide of the P-site codon to the 5' end of the FSS are colored coded and highlighted, respectively. The **sequences of the FSS variants and the biomolecular helices** are colored coded as in Figure 1. Every model mRNA contains a **handle sequence** complementary to the biotinylated DNA oligo, which is used to immobilize the mRNA in smFRET experiments.

**Supplementary table 2. The HIV and dnaX bimolecular helices formed by annealing two complementary RNA strands.**

| | ΔG (kcal/mol) | $K_{eq}$ | $k_{off}$ ($s^{-1}$) | $t_{1/2}$ (s) | [Input] (μM) | [Free RNA] (μM) | [bimolecular helix] (μM) |
|---|---|---|---|---|---|---|---|
| **HIV bimolecular helix (310.15 K)** | **-27.8** | **$3.90\times10^{19}$** | **$2.56\times10^{-14}$** | **$2.70\times10^{13}$** | **0.60** | **$1.24\times10^{-7}$** | **0.60** |
| **HIV bimolecular helix (293.15 K)** | **-34.8** | **$4.27\times10^{28}$** | **$2.34\times10^{-23}$** | **$2.96\times10^{22}$** | **0.30** | **$2.65\times10^{-12}$** | **0.30** |
| **dnaX bimolecular helix (310.15 K)** | **-28.6** | **$1.43\times10^{20}$** | **$7.00\times10^{-15}$** | **$9.90\times10^{13}$** | **0.60** | **$6.48\times10^{-8}$** | **0.60** |
| **dnaX bimolecular helix (293.15 K)** | **-39.7** | **$3.99\times10^{29}$** | **$2.51\times10^{-24}$** | **$7.66\times10^{19}$** | **0.30** | **$8.68\times10^{-13}$** | **0.30** |

In accordance with the incubation temperatures used in filter-binding and smFRET experiments, free energy change ΔG for RNA annealing were calculated at 37℃ (310.15 K) or at room temperature (293.15 K), respectively. Equilibrium constants $K_{eq}$ were calculated from $K_{eq}=\exp(-\Delta G/RT)$, where temperature T was either 310.15 or 293.15 K. $R=1.987\times10^{3}$ $kcal\cdot K^{-1}\cdot mol^{-1}$. Dissociation rates $k_{off}$ were calculated using equation $K_{eq}=k_{on}/k_{off}$, where association rate $k_{on}$ was approximated to $10^{6}$ $s^{-1}$ as its value is less temperature sensitive[4]. Half-life $t_{1/2}$ of each bimolecular helix were calculated from $t_{1/2}=\ln(2)/k_{off}$. In the calculation, prior to annealing, initial concentrations of the two complementary RNA strands [input] were equal (0.6 or 0.3 μM). Thus, concentrations of free and annealed RNAs, [free RNA] and [bimolecular helix], were determined by solving equations [bimolecular helix]+2×[free RNA]=2×[input] and $K_{eq}$=[bimolecular helix]/[free RNA]$^{2}$.
